# Pathway engineering in yeast for synthesizing the complex polyketide bikaverin

**DOI:** 10.1101/783365

**Authors:** Meng Zhao, Yu Zhao, Qi Hu, Hala Iqbal, Mingdong Yao, Hong Liu, Bin Qiao, Chun Li, Christine A. S. Skovbjerg, Jens Christian Nielsen, Jens B. Nielsen, Rasmus J.N. Frandsen, Yingjin Yuan, Jef D. Boeke

## Abstract

Fungal polyketides display remarkable structural diversity and bioactivity, and therefore the biosynthesis and engineering of this large class of molecules is therapeutically significant. Here, we successfully recoded, constructed and characterized the biosynthetic pathway of the formation of bikaverin, a tetracyclic polyketide with antibiotic, antifungal and anticancer properties, in *S. cerevisiae*. We used a green fluorescent protein (GFP) tagging strategy to identify the low expression of *Bik1* (polyketide synthase) as the bottleneck step in the pathway, and a promoter exchange strategy to increase expression of *Bik1* and bikaverin yield. To further increase product yield, we used an enzyme-fusion strategy to couple the monooxygenase (Bik2) and methyltransferase (Bik3) to efficiently channel intermediates between modifying enzymes, leading to a dramatic improvement of Bikaverin yield of nearly 60-fold. This study demonstrates that the biosynthesis of complex polyketides biosynthesis can be established and efficiently engineered in *S. cerevisiae*, highlighting the great potential for natural product synthesis and large-scale fermentation in yeast.

## Introduction

Fungal polyketides represent a large group of structurally and functionally diverse natural products with clinically-relevant medical properties, including antibiotic, antifungal, immunosuppression and anticancer activities^1–4^. This diversity is biosynthetically generated by the combination of modifying enzymes such as methyltransferases and oxygenases, and a multifunctional enzyme polyketide synthase (PKS) that condenses multiple carboxyl units in a highly-regulated, iterative process involving initiation, elongation, cyclization, and release^5^. Because the resulting molecules are often excellent candidates for novel therapeutics, the metabolic engineering of fungal polyketides, including both their biosynthesis and large-scale fermentation, is of substantial interest.

*Saccharomyces cerevisiae* has been one of the most widely used hosts for the metabolic engineering of natural products including antibiotics, terpenoids, cannabinoids and opiates^6–9^. Despite the genetic tractability and scalability of metabolic engineering in yeast, there are only a few examples of the successful biosynthesis of polyketides in *S. cerevisiae* and, to the best of our knowledge, no examples of further pathway engineering^10–14^. Because of their large enzyme size, complicated biosynthetic pathways and highly regulated processes, the biosynthesis and large-scale fermentation of polyketides in yeast remains a challenge.

To expand the metabolic engineering of polyketides in yeast, we constructed and characterized a cell factory for the production of the fungal polyketide bikaverin in *S. cerevisiae*. Bikaverin is a red-colored tetracyclic polyketide with antibacterial and anticancer activity produced by members of the genus *Fusarium*15-19. A putative bikaverin biosynthetic gene cluster has been identified in *Fusarium fujikuroi*, and a biosynthetic pathway has been proposed based on knockout-based studies, but the pathway has not been fully characterized nor verified via gain of function experiments. Among the 6 proteins encoded, enzymes Bik1, Bik2 and Bik3 are predicted to be responsible for the synthesis of bikaverin, while Bik4 and Bik5 are predicted transcription regulators, and Bik6 is a permease^18^. The polyketide carbon-skeleton of bikaverin is produced by the Type I PKS Bik1 (formerly called PKS4), which is a very large (over 2000 amino acid residues) multifunctional enzyme consists of the following domains: Starter unit acyltransferase (SAT), β-Ketosynthase (KS), Malonyl:ACP acyltransferase (MAT), Product template (PT), Acyl carrier protein (ACP), and Claisen cyclase (CLC)^20,21^. The SAT domain initially selects acetyl-CoA as a starting unit, and the KS and MAT domains condense eight units of malonyl-CoA to the growing polyketide chain, resulting in a nonaketide chain which is covalently tethered to the ACP domain. The PT and TE/CLC domains cyclize the chain, releasing the intermediate pre-bikaverin from Bik1. Pre-bikaverin is modified with two oxidations catalyzed by the FAD-dependent monooxygenase Bik2, and two methylations catalyzed by *O*-methyltransferase Bik3, to form the final product bikaverin (Fig. 1a).

**Fig. 1.**
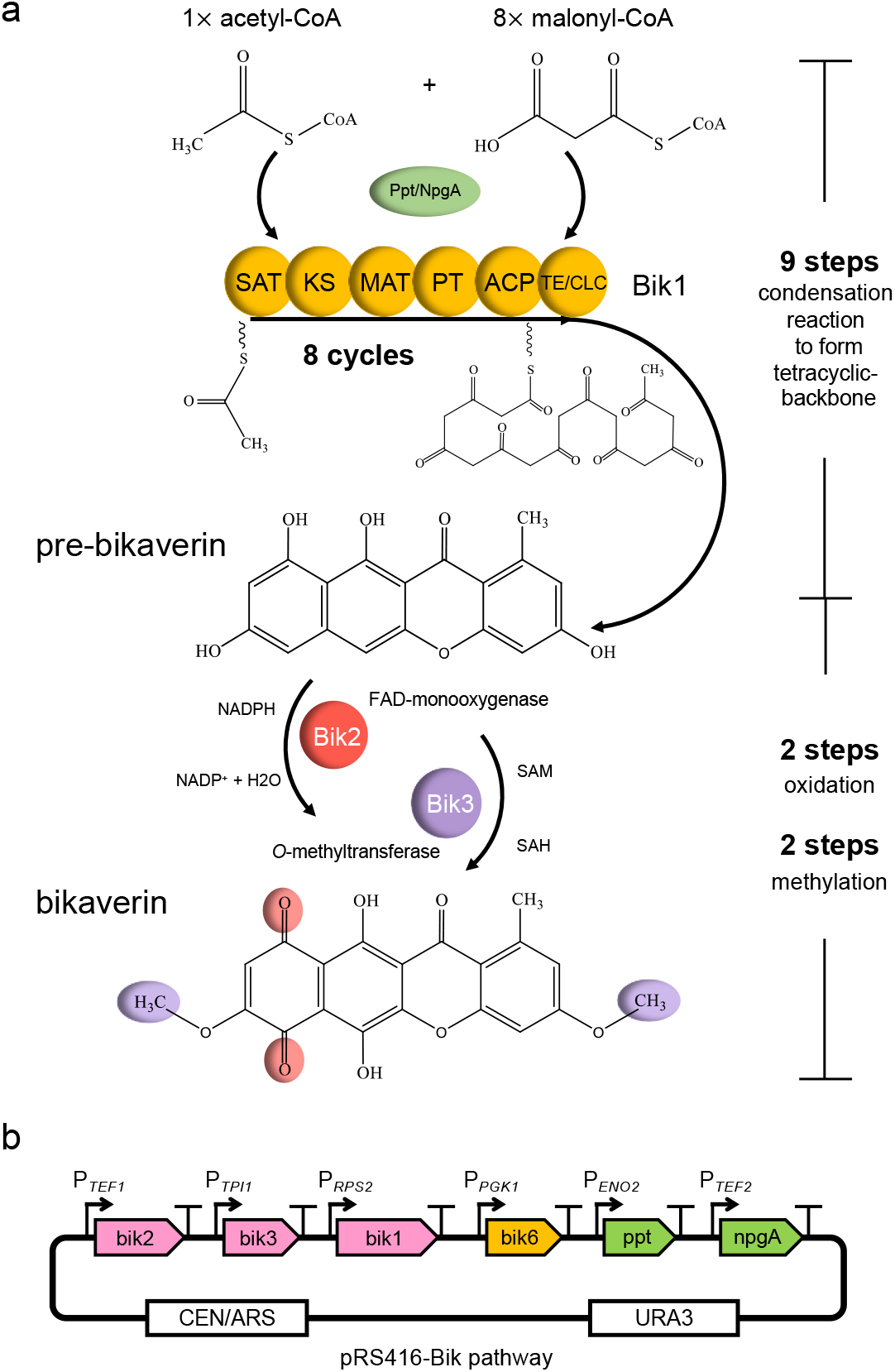
Bikaverin biosynthetic pathway in *S. cerevisiae*. **a** In the bikaverin biosynthetic pathway, the polyketide synthase Bik1, activated by phosphopantetheinyl transferase (PPTase), condenses one acetyl-CoA and eight malonyl-CoA units to form pre-bikaverin. The modifying enzymes Bik2 (monooxygenase) and Bik3 (*O*-methyl-transferase) convert pre-bikaverin into the final product bikaverin. **b** The constructed bikaverin pathway encoding plasmid containing *ppt1*, *npgA* and two versions of bik genes (*bik1*, *bik2*, *bik3*, *bik6*) was codon-optimized, synthesized and assembled into pRS416 plasmid for transformation into *S. cerevisiae*.

In this study, we successfully recoded, constructed and characterized the biosynthetic pathway of the complex polyketide bikaverin in *S. cerevisiae*. We developed a GFP-tagging method for identifying the expression of *bik1* as the bottleneck in the pathway and used a promoter exchange strategy to increase expression of *bik1* and subsequently, bikaverin yield. In addition, we verified the genes essential for bikaverin synthesis and characterized the biosynthetic pathway through a bottom-up strategy. Finally, we optimized bikaverin production by developing an enzyme-fusion method to couple the modifying enzymes Bik2 and Bik3, allowing channeling of intermediates between the enzymes and achieving a ~60-fold increase in the final yield of the molecule. The three strategies we describe expand the toolbox of polyketide biosynthesis in *S. cerevisiae*, and will enable development of yeast as a host for the biosynthetic engineering of fungal polyketides and other desired molecules.

## Results

### Establishing the biosynthesis of bikaverin in *S. cerevisiae*

To heterologous express the bikaverin biosynthetic pathway in *S. cerevisiae*, we designed a yeast-recoded bikaverin gene cluster comprising the genes *bik1*, *bik2*, *bik3* and *bik6*, as well as the PPTase (phosphopantetheinyl transferase) needed for post-translational activation of the ACP domain in the PKS enzyme. We retrieved the protein sequences of Bik1, Bik2, Bik3, and Bik6 from two *F. fujikuroi* records in the NCBI Protein database: version 1 (accession numbers: Bik1, CAB92399; Bik2, CAJ76275; Bik3, CAJ76274; Bik6, CAM90596) from 2017 and a revised version 2 (accession numbers: Bik1, S0DZM7; Bik2, S0E2X6; Bik3, S0E608; Bik6, S0DZN4) from 2013^22^. Most of the protein sequences were identical in the two versions, excepting a few single amino acid variants and gaps (Supplementary Fig. 1). We synthesized *S. cerevisiae*-optimized *bik1*, *bik2*, *bik3* and *bik6* genes from both versions (V1-2007 and V2-2013) under the control of the yeast endogenous promoters, P*RPS2*, P*TEF1*, P*TPI1* and P*PGK1*, respectively (Fig. 1b). In addition, to facilitate the post-translational activation of the PKS, we chose two PPTase’s to activate Bik1: Ppt1, the native PPTase from *F. fujikuroi*, and NpgA, a PPTase from *A. nidulans*, which previously has been used successfully in yeast for activating PKSs and nonribosomal peptide synthetases (NPRSs)^6,12,13^. Both enzyme coding genes were added into the constructed yeast bikaverin expression cassette under the control of the *ENO2* and *TEF* promoter, respectively. The resulting bikaverin expression cassette was transformed into yeast, as strain yZM001 for V1-2007 and yZM006 for V2-2013. The strains failed to turn red, suggesting that the red-colored compound bikaverin was not being produced.

To probe the bikaverin pathway for bottlenecks limiting biosynthesis, we developed a GFP-fusion strategy to confirm that all genes were well expressed and translated into proteins efficiently. The coding sequence for green fluorescent protein (GFP) was fused, in frame, to the 3’ end of the individual genes. When the genes *bik2*, *bik3*, *bik6*, *ppt* and *npgA* were tagged, the cells displayed a strong fluorescent signal, whereas cells meant to express *bik1* displayed a weak signal suggesting that the Bik1 level was limiting bikaverin biosynthesis (Fig. 2a). This may be due to the very large size of *bik1* relative to the other genes, or due to toxicity resulting from *bik1* expression, and subsequent selection for non-expressing cells. To optimize *bik1* expression, we used a promoter exchange strategy to replace the initially used promoter P*RPS2* with several different yeast endogenous promoters: P*RPL43A*, P*GPM1*, and P*GAL1*. The Bik1-GFP fusion was strongly expressed only when driven by the galactose-inducible *GAL1* promoter (strain yZM009, using V2-2013 enzymes) (Fig. 2b). Consistent with this result, strain yZM009 only turned red when grown on galactose, indicating that bikaverin was being produced (Fig. 2c). Production of the compound was confirmed using HPLC (Fig. 2d) and MS (Supplementary Fig. 2).

**Fig. 2.**
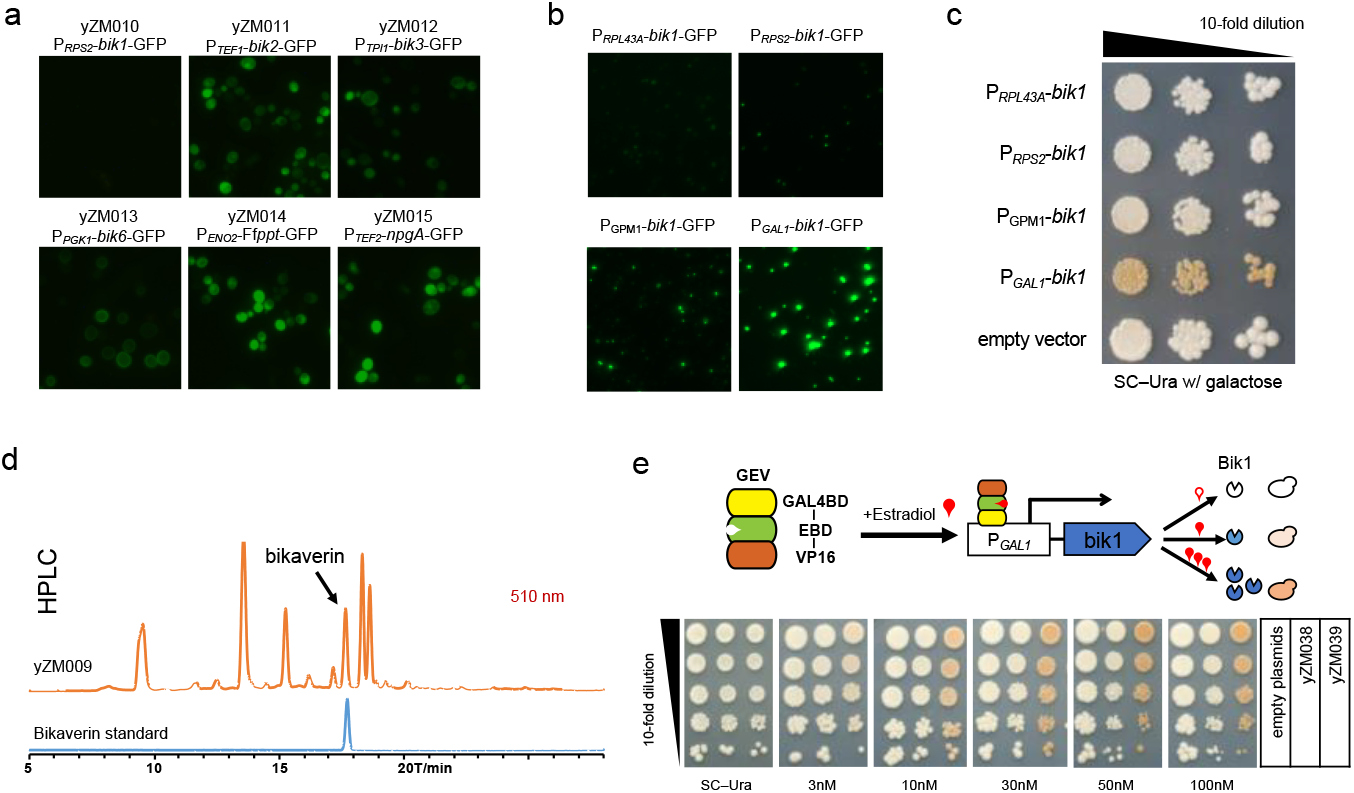
GFP-Fusion strategy for identifying bottlenecks in the bikaverin pathway. **a** Bikaverin pathway genes were tagged with GFP to check expression and identify *bik1* as the bottleneck in the pathway. **b** Promoter exchange strategy was used to identify P*GAL1* as driving high expression of *bik1.* **c** Dot assay showing that colonies with the P*GAL1*-*bik1* turned red, indicating the presence of bikaverin. **d** HPLC analysis confirmed the production of bikaverin by strain yZM009. **e** Inducible system to regulate P*GAL1*-*bik1* expression with estradiol instead of galactose verified that bikaverin was produced via the promoter exchange. After two days’ incubation, yeast colonies displayed deeper color when grown on higher β-estradiol concentration in glucose medium.

To confirm that bikaverin biosynthesis was due to higher expression of *bik1* and not because of the use of galactose instead of glucose as the carbon source, we introduced a galactose-independent system to regulate the GAL1 promoter. We expressed a GAL4(1-93).ER.VP16(GEV) plasmid to produce a tribrid protein comprising a GAL4 DNA binding domain, an estrogen receptor and a VP16 domain^23^. When bound to β-estradiol, this complex translocate into the nucleus and binds to P*GAL1* sequences to activate gene transcription; consequently, an increase in β-estradiol availability leads to higher rates of transcription^23^. With *bik1* under the control of this system, increasing the concentration of β-estradiol in the medium led to deeper red colony color (Fig. 2e), indicating higher yields of bikaverin and confirming that the higher expression of *bik1* indeed resulted from the promoter exchange strategy.

We also observed that the V1-2007 strain yZM002 did not turn red using the same promoters as used in yZM009, and no bikaverin was produced. We further checked the function of each V1-2007 enzyme in yeast by replacing them with their counter part from the V2, and found all three V1 proteins were expressed much less efficiently (Supplementary Fig. 3). As a result, V2-2013 clusters were used for further characterization.

### Characterization of the bikaverin pathway in *S. cerevisiae*

With the production of bikaverin successfully established in *S. cerevisiae*, we turned to the characterization of the biosynthetic pathway using both top-down and bottom-up strategies. To answer questions about the essentiality of *bik6* for bikaverin production in yeast and whether both PPTases can activate Bik1, we used a top-down approach to delete *bik6*, *ppt1* and *npgA* separately and in combination from the yeast-recoded bikaverin pathway. Unlike in *F. fujikoroi*, in which deletion of *bik6* has been reported to reduces bikaverin yield, deletion of *bik6* in *S. cerevisiae* had minimal effect on bikaverin yield (Fig. 3a). Deletion of either *ppt1* or *npgA* reduced bikaverin yield 2-fold compared to the original strain. However, bikaverin was still successfully synthesized in both cases, indicating that both Ppt1 and NpgA are capable of activating the ACP domain of Bik1, but that one copy of either gene did not provide enough catalytic potential to allow for post-translational activation of the produced Bik1 enzyme.

**Fig. 3.**
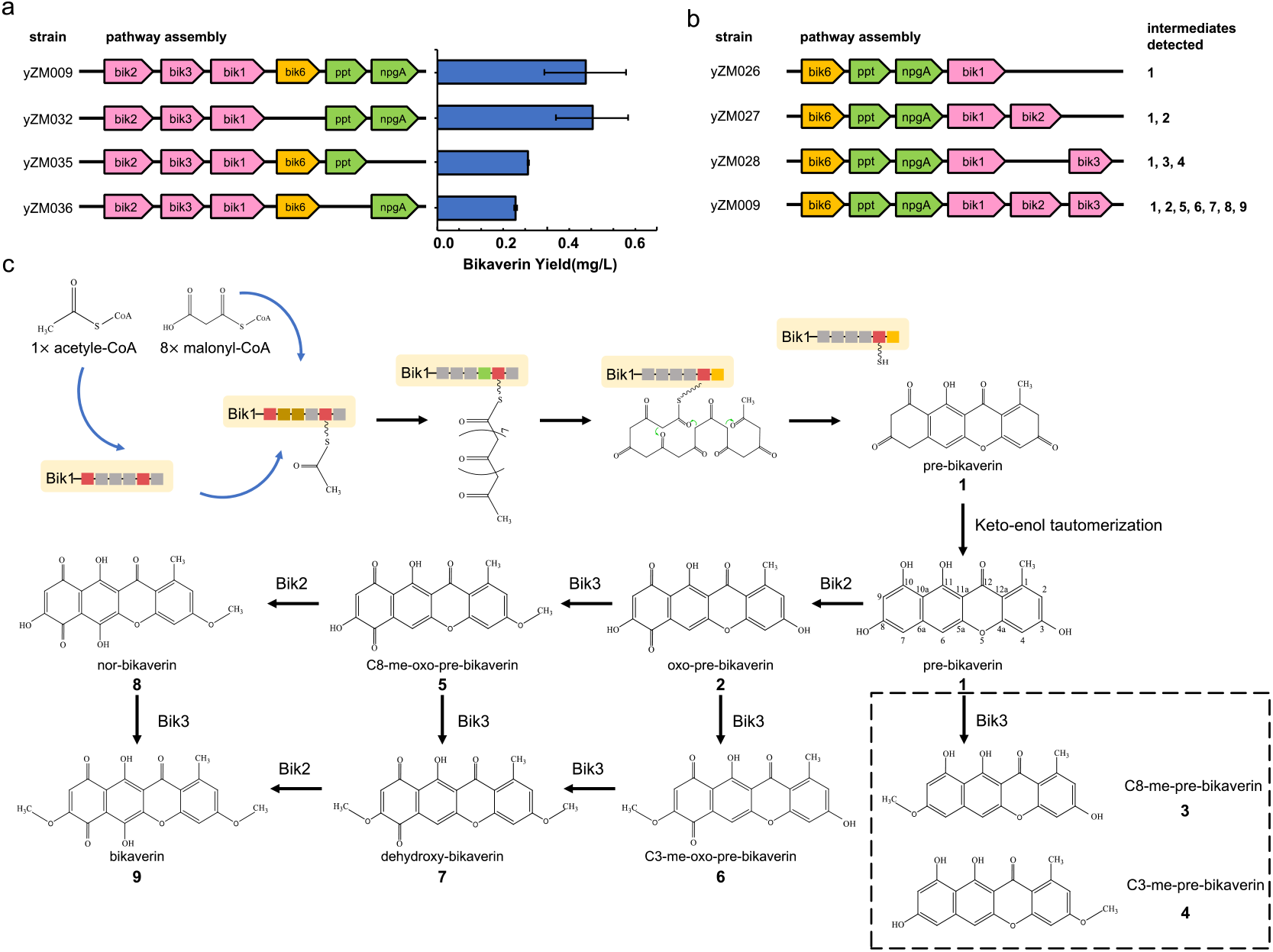
Characterization of bikaverin synthesis in yeast. **a** A top-down strategy was used to measure the effects of *bik6*, *ppt* and/or *npgA* deletion on bikaverin production. **b** A bottom-up strategy was used to build constructs with different combinations of *bik1*, *bik2* and *bik3* genes. The identification of resulting intermediates was detected by HPLC-MS (Supplementary Fig. 5-8) and used to decipher the biosynthetic pathway of bikaverin. **c** The putative bikaverin biosynthesis pathway in *S. cerevisiae*. Boxed intermediates were only detected in the absence of *bik2* (Supplementary Fig. 6).

To further characterize bikaverin biosynthesis and to establish the order of reactions in the pathway, we used a bottom-up strategy to build constructs with different combinations of *bik1*, *bik2* and *bik3* and analyzed the resulting strains using HPLC-MS^24^. As expected, pre-bikaverin (m/z 325.06) (**1**) was detected in all strains containing *bik1* (Fig. 4b, c and Supplementary Fig. 5). Oxo-pre-bikaverin (m/z 339.04) (**2**), the expected intermediate resulting from the C7 oxidation of pre-bikaverin by the monooxygenase Bik2, was detected in strains containing *bik2* (yZM027 and yZM009). Interestingly, di-nor-bikaverin, the predicted product from the further oxidation of C7-oxo-pre-bikaverin, was not detected in either strain expressing Bik2 (yZM027 and yZM009), indicating that the monooxygenase cannot oxidize the C6 position without prior oxidation of the C7 positions (Supplementary Fig. 6).

**Fig. 4.**
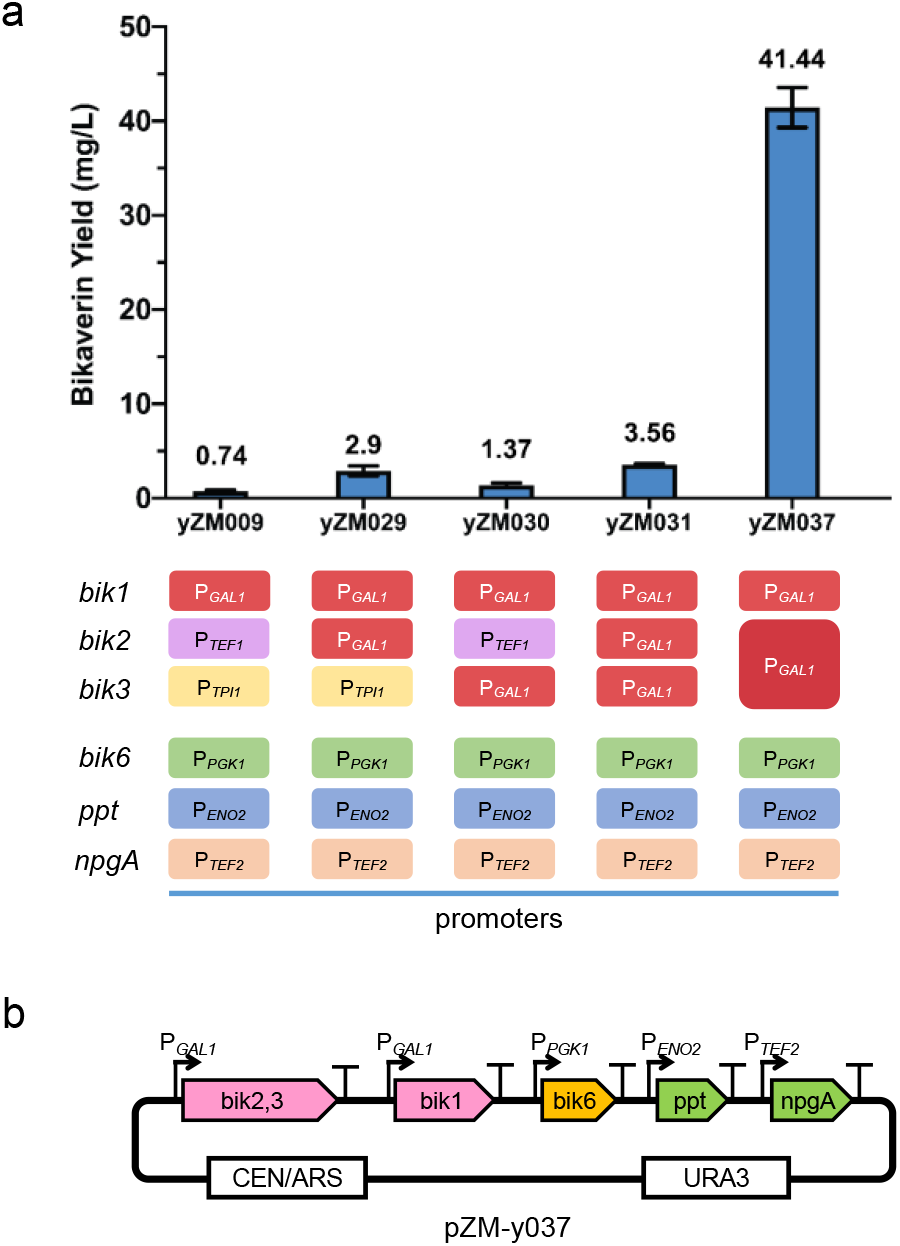
Optimization of bikaverin yield using promoter exchange and enzyme fusion strategies. **a** Driving *bik2* and *bik3* using GAL1 lead to modest increases in bikaverin yield. Fusing bik2 and bik3 led to a dramatic increase in bikaverin production, 60-fold higher than the original strain. **b** The bikaverin pathway construct containing *bik2* and *bik3* fused to form the *bik2,3* fusion gene. The whole pathway was assembled into a plasmid vector with *URA3*.

In the strain containing only *bik1* and *bik3* (yZM028), two small peaks of m/z 339.09 (**3**) and 339.08 (**4**) were detected (Supplementary Fig. 7). HPLC-MS retention time confirmed that neither peak was oxo-pre-bikaverin, and neither peak was detected in strains that lacking *bik3*. We speculated that these peaks are intermediates resulting from methylation of pre-bikaverin to form C8-me-pre-bikaverin (**3**) and C3-met-pre-bikaverin, catalyzed by Bik3. The two peaks were small and not detected in strain yZM009 containing the entire pathway, suggesting that pre-bikaverin can be methylated by Bik3 but that this is unfavored relative to oxo-pre-bikaverin and other downstream substrates including C3/C8 me-oxo-pre-bikaverin and nor-bikaverin.

In strain yZM009 containing all the pathway genes, two peaks of m/z 353.05 were detected (Supplementary Fig. 8c). These were postulated to be the two mono-methylated forms of oxo-pre-bikaverin: C3-me-oxo-pre-bikaverin (**6**) and C8-me-oxo-pre-bikaverin (**5**). In addition, a peak of dehydroxy-bikaverin (m/z 367.07) (**7**), the di-methylated form of oxo-pre-bikaverin, was detected in this strain (Supplementary Fig. 8d), suggesting that Bik3 is able to methylate oxo-pre-bikaverin twice in yeast. Nor-bikaverin (**8**), the intermediate produced from the further oxidation of C8-me-oxo-pre-bikaverin by Bik2, was also detected (m/z 369.05) and confirmed by comparison to the previously published MS trace of the molecule^24^. However, at m/z 369.05, only one peak was detected, which suggested that C8-me-oxo-pre-bikaverin, not C3-me-oxo-pre-bikaverin, was the favored substrate of *bik2*, although it is possible that the oxidized form of C3-me-oxo-pre-bikaverin cannot be isolated using the HPLC-MS conditions tested (Supplementary Fig. 8e). With this certain preference for substrates, Bik2 only catalyzes the oxidation C8-me-oxo-pre-bikaverin, while Bik3 can methylate both C3- and C8-me-oxo-pre-bikaverin. The final product, bikaverin (m/z 383.06) (**9**), was only detected in the strain containing the complete pathway, confirming that Bik1, Bik2 and Bik3 are all essential and sufficient for the *de novo* biosynthesis of bikaverin (Supplementary Fig. 8f).

Using the results from the HPLC-MS experiments, we traced out the putative bikaverin biosynthetic pathway in yeast (Fig. 3c). Pre-bikaverin is first oxidized to oxo-pre-bikaverin by Bik2, then to dehydroxy-bikaverin by two methylation steps catalyzed by Bik3. Dehydroxy-bikaverin can then be oxidized to be form the final product bikaverin. In addition, the Bik2 intermediate C8-me-oxo-pre-bikaverin, but not C3-me-oxo-pre-bikaverin, can also be oxidized to nor-bikaverin and then methylated to form the final product bikaverin.

### Pathway optimization to increase Bikaverin production

To further optimize the yield of bikaverin, we applied the promoter exchange strategy to the modifying enzymes in the bikaverin pathway. The GFP-fusion strategy revealed that the expression of Bik2 and Bik3 was significantly lower than the expression of Bik1 following the use of the GAL1 promoter (Supplementary Fig. 9). Changing the promoter of *bik2* and *bik3* to P*GAL1* resulted in a 3-fold increase in bikaverin yield from ~0.74mg/L to ~2.9mg/L for *bik2* and an increase to ~1.37mg/L for *bik3* (Fig. 4a). Further, changing the promoter of both *bik2* and *bik3* to P*GAL1* resulted in a bikaverin yield of ~3.65mg/L, which was five-fold higher than the original strain yZM009.

Characterization of the bikaverin biosynthetic pathway highlighted that pre-bikaverin is modified in a stepwise manner by the monooxygenase Bik2 and then the *O*-methyltransferase Bik3. This process results in many intermediates, some of which are dead-end products. This led us to hypothesize that channeling the mono-oxygenated pre-bikaverin from Bik2 to Bik3 may increase the yield of bikaverin. Guided by homology modeling of the two enzymes, which revealed that the active pocket and FAD binding site of Bik2 are near its N-terminal and the SAM binding site and active pocket of Bik3 are close to C-terminal (Supplementary Fig. 10), we fused Bik2 and Bik3 together using a (GGGS)^3^ linker to bring the N-terminal of Bik2 and the C-terminal of Bik3 in close proximity^25^. The resulting strain pZM-y037 displayed a significant increase in the yield of bikaverin to ~41 mg/L, approximately 11-fold higher than the strain with P*GAL1* driving *bik1*, *bik2* and *bik3* (yZM031), and nearly 60-fold higher than the original strain producing bikaverin (yZM009) (Fig. 4a, b). The protein level of the Bik2-Bik3 fusion protein was not higher than the levels detected of Bik2 or Bik3 respectively when expressed as individual genes (Supplementary Fig. 11), indicating that the fusion protein improved bikaverin yield by increasing the efficiency of polyketide modification. This suggested that building an intermediate channel for staged catalytic reactions can significantly increase biosynthesis efficiency.

## Discussion

Fungal polyketides display remarkable structural and functional diversity and are excellent candidates for the development and biosynthetic engineering of new therapeutics with anticancer, antifungal, antibiotic and immunomodulation properties. In this study, we demonstrated the efficient biosynthesis of the complex fungal polyketide bikaverin in the heterologous host *S. cerevisiae*. To achieve this, we reconstructed the bikaverin pathway in yeast by recoding and synthesizing the genes *bik1*, *bik2*, *bik3*, and *bik6* from *F. fujikoroi* along with two PPTases that activate the ACP domain of the PKS Bik1.

To optimize production of bikaverin, we used three strategies to debug the biosynthetic pathway and increase the finale yield, and these can be extrapolated to the production of other small molecules in yeast. We used a GFP-fusion strategy to quickly identify low Bik1 expression as the initial bottleneck step in the bikaverin biosynthetic pathway (Fig. 5a). We used a promoter exchange strategy to increase expression of Bik1, and subsequently increase bikaverin yield (Fig. 5b). This strategy was also used to increase expression of the modifying enzymes Bik2 and Bik3, further increasing the yield of the final product. In addition, we used an enzyme-fusion strategy to fuse together the modifying enzymes Bik2 and Bik3 to efficiently channel intermediates through the pathway, leading to an 11-fold increase in bikaverin yield (Fig. 5c). We postulate that this fusion may help to channel substrates from one modifying enzyme to the other, preventing the diffusion of intermediates and improving the efficiency of substrate transfer between the two enzymes. In many biosynthetic pathways, modifications such as the methylation, glycosylation and hydroxylation occur step-wise^26–29^; this protein fusion strategy might similarly be applied to such pathways for improving yield.

**Fig. 5.**
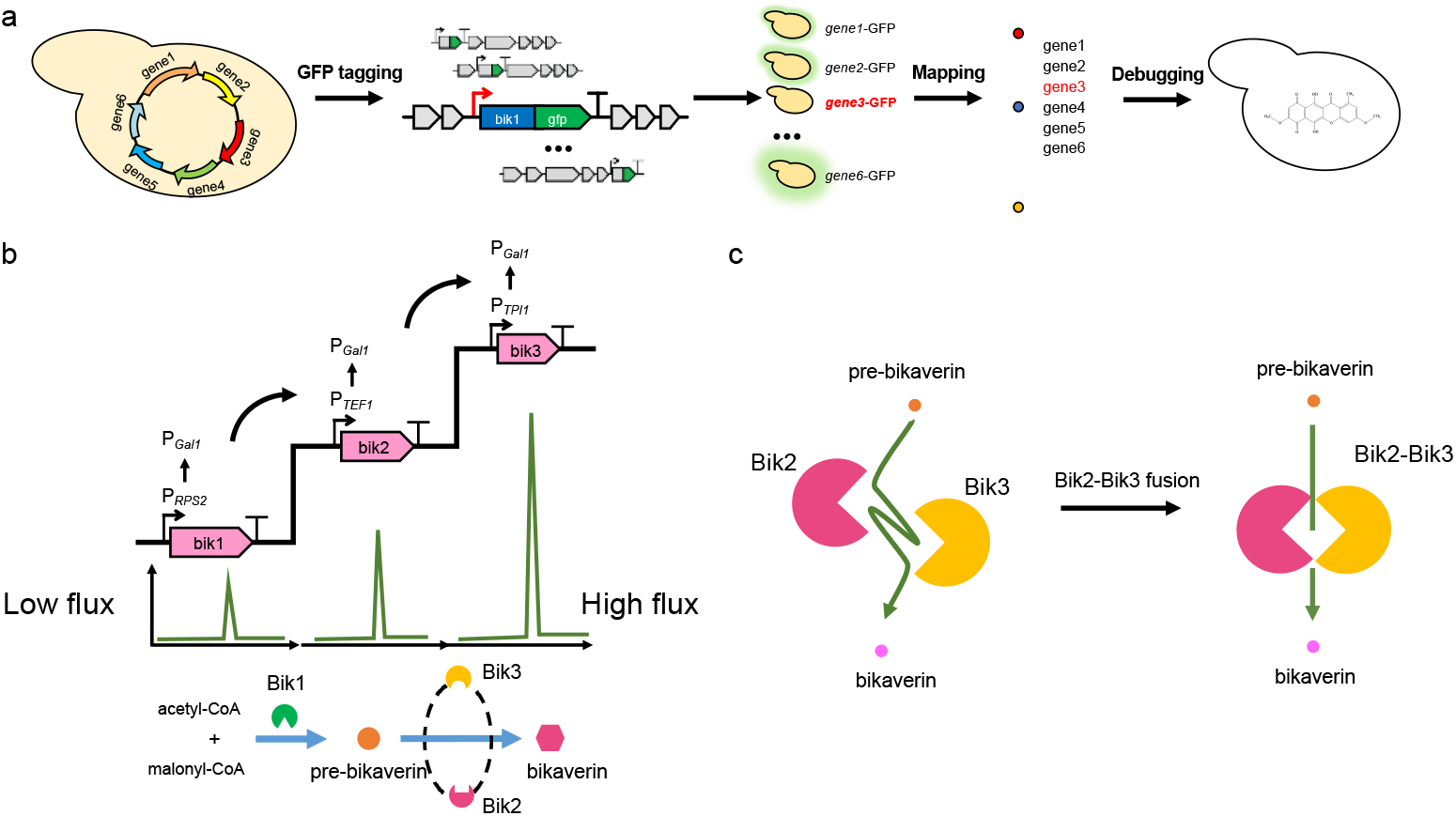
Strategies for optimizing bikaverin synthesis in *S. cerevisiae*. **a** GFP-Fusion strategy to debug gene expression. **b** Promoter exchange strategy to increase expression of key biosynthetic genes in the pathway. **c** Channeling substrates by enzyme fusion. Enzyme-fusion strategy was used to stage the pathway to increase yield by channeling intermediates between the modifying enzymes.

Previously, bikaverin was biosynthesized in its native host *F. fujikoroi*^18^. Yeast is a significantly better model organism for the biosynthetic engineering of fungal polyketides, with many available synthetic biological tools including pathway assembly, CRISPR genome editing and even whole chromosome synthesis and SCRaMbLE ^30–36^. Using the synthetic biology tools available in yeast, we were able to characterize the bikaverin biosynthetic pathway using top-down and bottom-up strategies. Our result showed that genes *bik1*, *bik2* and *bik3* are essential for the production of bikaverin, while omitting of the permease encoding gene *bik6* had minimal effect in yeast. We were also able to confirm the production of the proposed intermediates of the bikaverin biosynthetic pathway and determine the order of the modification steps catalyzed by Bik2 and Bik3. In addition, the use of yeast promoters and terminators to drive expression of the bikavarin pathway genes led us to establish expression that was not dependent on the transcription regulation systems present in *F. fujikuroi.* Consequently, our strains display stable, tunable and scalable bikaverin production that is not dependent on environmental conditions. This is an added advantage of using yeast as a host for the bioengineering of polyketides.

The approach described in this study greatly expands the toolbox for biosynthetic engineering of fungal polyketide pathways in yeast. Moreover, in the polyketides biosynthesis, the type I iterative PKS enzyme works as an assembly line with each domain catalyzing a different function and this predictability of function implied that PKSs can be rationally engineered^5,37^. It has been reported that swapping of domains between PKSs results in new diverse functions *in vitro*38-40. The present results now allow for the implementation of similar combinatory hybrid PKS experiments in *S. cerevisiae* with the aim of create novel PKS enzymes that generate new polyketides. We hope that our study including the tools described in this study, as well as the suite of pre-existing synthetic biology tools, will promote biosynthetic engineering of PKSs in *S. cerevisiae*, allowing for the exploration of the chemical space of polyketides and the development of novel therapeutics with diverse bioactivities.

## Methods

### Yeast strains and plasmid assembly

All strains, plasmids used in this study were listed in Supplementary Table 1 and 2. The protein sequences of Bik1, Bik2, Bik3, Bik6, FfPpt1, NpgA were obtained from NCBI database; accession numbers were listed in Supplementary Figure 1. Protein sequences were reverse-translated into nucleotide sequences and codon optimized for expression in yeast using BioPartsBuilder^41^. Yeast promoters and terminators were PCR amplified from BY4742 genomic DNA. All pathway genes (*bik1*, *bik2*, *bik3*, *bik6*, *ppt1*, *npgA*) were individually constructed as transcription units (TUs) consisting of promoter, coding sequence and terminator employing the yeast Golden Gate (yGG) DNA assembly method^42^. We used the versatile genetic assembly system (VEGAS) to coassemble all the TUs into the pRS416 plasmid backbone in the BY4742 strain (*MAT*α *ura3∆0 leu1∆0 lys2∆0 his3∆1*)^43,44^. The final construct was miniprepped from yeast and transformed into *E. coli* for sequencing verification and storage. The correct plasmid with the complete bikaverin pathway was re-transformed back into BY4742 for further study.

Yeast strains were grown in YPD medium or SC dropout plates supplemented with appropriate amino acids and/or drugs. All yeast transformations in this study were done using standard lithium acetate protocols^44^.

### Fusion protein construction in GFP-Fusion strategy

To add a GFP tag to the C-terminal of each coding sequence, the GFP-KanMX6 fragments were PCR amplified with corresponding homologous arms. Then the PCR amplicon was transformed into the yeast strain carrying the complete bikaverin pathway. Successful integration was selected on YPD+G418 plate and confirmed by colony PCR and Sanger sequencing.

### Fluorescence microscopy and intensity measurement

Pictures were taken using an EVOS-FL Auto cell imaging system (Invitrogen) with the 20X objective lens. Yeast cells were cultured on plates using appropriate selective media for 2 days, then picked onto wet mount slides for visualization. To measure relative GFP fluorescence intensity, single colonies were picked into 24 deep-well plates with 1.5ml SC–Ura medium in each well, then incubated at 30°C, 250 rpm/min for 20 hours. Cultures were diluted (1:20) in nonopaque 96 well plates to measure A_600_ and opaque 96 well plates to detect GFP fluorescence intensity. In this study, relative fluorescence intensity was defined as GFP fluorescence intensity / A_600_.

### Strain fermentation and bikaverin extraction

Yeast strains were cultured in 5 ml of SC–Ura liquid medium at 30°C, 250 rpm, overnight to saturation. Cells were harvested by centrifugation at 5000 rpm for 5 min, followed by 3x 2 ml washes with water, and then inoculated to an initial OD_600_ of 0.5 in 30 ml of SC–Ura liquid medium with either glucose or galactose as the carbon source. Cultures were incubated at 30°C, 250 rpm, for 96 hours. Appropriate volumes of cell cultures (5-2 ml) were centrifuged for 10 min at 12000 rpm. Cell pellets were resuspended with 0.5 ml water in a 2 mL tube, and 200 uL of acid-washed glass beads (Sigma, G8772-100G) were added into the tubes along with 0.5 mL ethyl acetate. Resulting suspensions were vortexed for 20 min and centrifuged for 10 min at 12,000 rpm before the supernatant was transferred to a new tube; this was repeated 5 times. Finally, extracts were dissolved in 500 µL acetonitrile (ACN) and centrifuged 3 times at 12000 rpm, 10 min. Resulting extracts were used for HPLC and HPLC-MS analysis.

### HPLC analysis

HPLC analysis of bikaverin was performed on a Water 2695 system equipped with a 2489 UV detector operating at 510 nm. Separation was achieved using a HyPURITY C18 5um 150×4.6mm column (Thermo Fisher, 22105-154630) at 25°C. The HPLC program was as follows: ACN as solvent A and 0.1% formic acid as solvent B; flow rate 0.8 ml/min; injection volume 20 µl; 0 min 30%A/70%B, 3 min 30%A/70%B, 5 min 35%A/65%B, 20 min 65%A/35%B, 25 min 80%A/20%B, 30 min 80%A/20%B, 30.5 min 30%A/70%B, 45 min 30%A/70%B. Bikaverin standards (Sigma, SML0724) at the following concentrations: 60 µg/ml, 30 µg/ml, 15 µg/ml, 7.5 µg/ml, 3.75 µg/ml, 1.87 µg/ml, were used to establish a standard curve to calculate yields.

### LC/ESI-MS analysis

All MS data was measured by a HPLC/ESI-MS method. Bruker micrOTOF-Q II instrument connected with Agilent HPLC system (Agilent G1312B SL binary; Agilent G1367C SL WP) were used to detect bikaverin and pathway intermediates. The HPLC program and column using in this experiment were same as the method in HPLC analysis. Electrospray ionization (ESI) mass spectrometry was performed in positive mode and set as follows: voltage set capillary at 4500V, set end plate offset at −500V; nebulizer 2.0 Bar; Dry heater 180°C; Dry Gas 6.0 l/min. The Mass date was analyzed using Compass Data Analysis software.

### Homology modeling and structural analysis in silico

To explore the effect of protein fusion version of Bik2 and Bik3, the tridimensional structure models of Bik2, Bik3 and forward fusion-protein Bik2-Bik3 were built and optimized by EasyModeller^45^. The high-resolution complex structures of rifampicin monooxygenase with FAD (PBDID: 5KOW) and methyltransferase with SAM (PBDID: 5w7p-A) were used as the templates of Bik2 and Bik3, respectively.

Modeled structures were analyzed using Pymol software^46^.

## Acknowledgements

We thank Prof. Gungrong Zhao and Yunzi Luo for their helpful discussions during manuscript preparation. This work was supported in part by National Natural Science Foundation of China (21621004, 21750001, 31861143017) to Y.J.Y. This work was also supported in part by NSF (MCB-1616111) to J.D.B. and in part by the Novo Nordisk Foundation (NNF15OC0016626) to R.J.N.F.

## Author contributions

J.C.F.N, J.B.N., and J.D.B. conceived the project. M.Z. performed all major experiments. Q.H. contributed to strain constructions and sample preparations for HPLC/MS. M.D.Y. did fusion protein homology modeling. M.Z. and H.L. performed LCMS analysis. B.Q. performed MS analysis. C.A.E.S. performed HPLC-MS analyses supervised by R.J.N.F.. J.C.F.N. and R.J.N.F contributed to initial pathway design. M.Z. and Y.Z. analyzed data and prepared figures. M.Z., Y.Z., H.I., Y.J.Y., and J.D.B., wrote the manuscript and all authors edited the manuscript. This project was supervised by Y.J.Y. and J.D.B.

