## Supplementary Figures for "Pathway engineering in yeast for synthesizing the complex polyketide bikaverin"

This supplementary figure ppt file includes:

**Supplementary Fig. 1**. Sequence alignment result between two versions of bik genes.

**Supplementary Fig. 2.** More HPLC and UV spectra detection of bikaverin in yZM009.

**Supplementary Fig. 3.** Bikaverin production from V1 and V2 chimeric pathway.

**Supplementary Fig. 4.** The Bikaverin standard for HPLC/ESI-MS analysis.

**Supplementary Fig. 5.** The EIC of pre-bikaberin with m/z 325.1 in strain yZM026, 027, 009.

**Supplementary Fig. 6**. The EIC of oxo-pre-bikaberin and dinor-Bikaverin in strain yZM027.

**Supplementary Fig. 7.** The EIC of me-pre-bikaberin from strain yZM028.

**Supplementary Fig. 8.** HPLC/ESI-MS spectrum of  strain yZM009, with the complete bikaverin pathway.

**Supplementary Fig. 9.** Relative expression of bik genes.

**Supplementary Fig. 10.** Bik2-Bik3 fusion design guided by homology modeling.

**Supplementary Fig. 11.** The expression level of Bik2-Bik3 fusion protein, compared to Bik2 and Bik3 expressed separately.

| Name | Protein accession | Alignment |
|---|---|---|
| Bik1 | V1: CAB92399<br>V2: S0DZM7 | V1 95 IDRC---------------------------------------------------CHCA 102<br>V2 95 IDYLARSDKQHPPAAPSLLLGICTGSIAAAAVSCA 129<br>V1 534 NTAIAGGTNVMTNPDNFAGLDRGHFLSRTGNCKAFN 569<br>V2 561 DTAIAGGTNVMTNPDNFAGLDRGHFLSRTGNCKAFD 596 |
| Bik2 | V1: CAJ76275<br>V2: S0E2X6 | V1 165 VRTLVRTHIDSKLPEPLTADDLHQRCLLHSLRHVSTHRSIA 205<br>V2 165 VRSAVRTHIDSKLPEPLTADDYISVACSTVYGMSAPTEGIA 205 |
| Bik3 | V1: CAJ76274<br>V2: S0E608 | V1 144 GVLKLR 149<br>V2 144 GVAELR 149 |
| Bik6 | V1: CAM90596<br>V2: S0DZN4 | T55I, S122A, S233G,F319L |

**Supplementary Fig. 1**. Sequence alignment result between two versions of bik genes. The differences in amino acid sequence were listed in alignment line. Their protein accession numbers are from NCBI Protein database (https://www.ncbi.nlm.nih.gov/protein).

### HPLC

yZM009

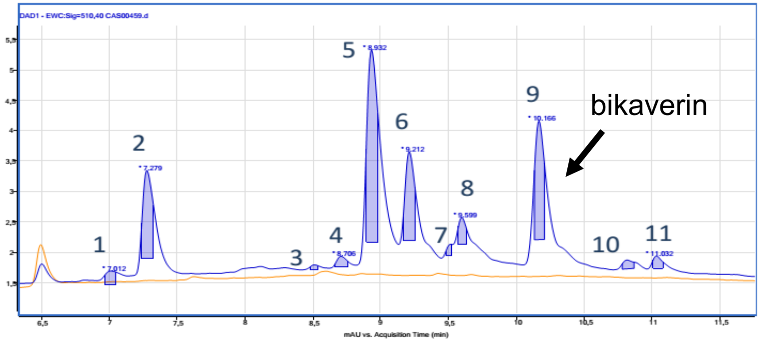


**Supplementary Fig. 2**. More HPLC and UV spectra detection of bikaverin in yZM009. This was also confirmed by mass spectra analysis.

### UV spectra

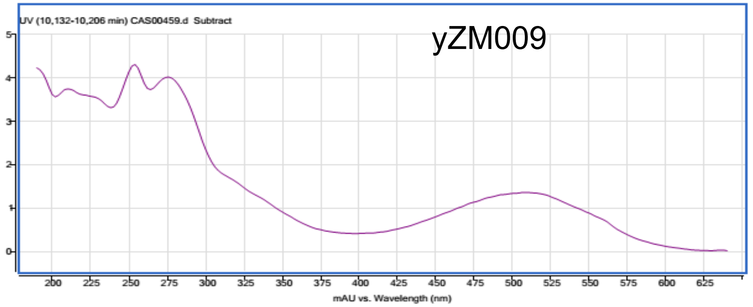


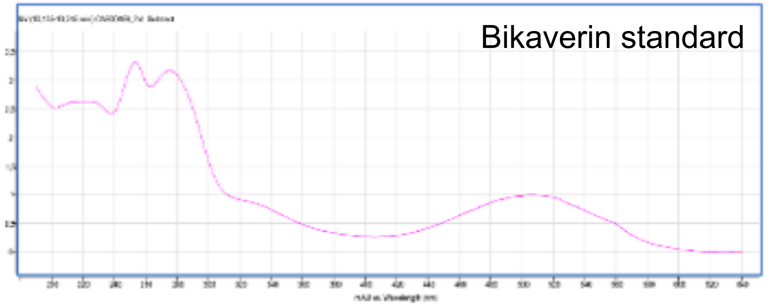


### Mass spectra

(Diagnostic adducts shown [M+H]+, [M+Na]+ and [2M+Na]+ )

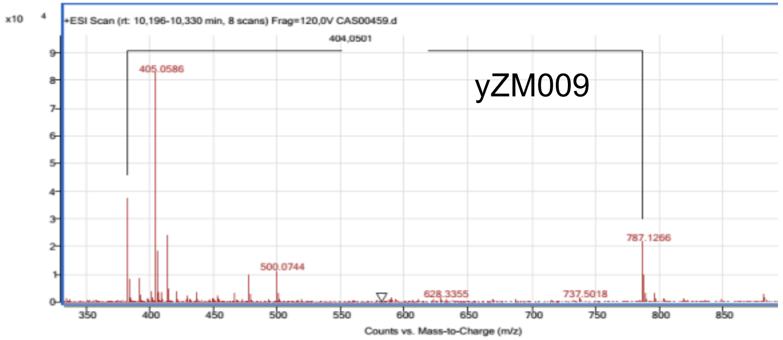


(Diagnostic adducts shown [M+H]+, [M+Na]+. [2M+Na]+ was also found but not shown on figure)

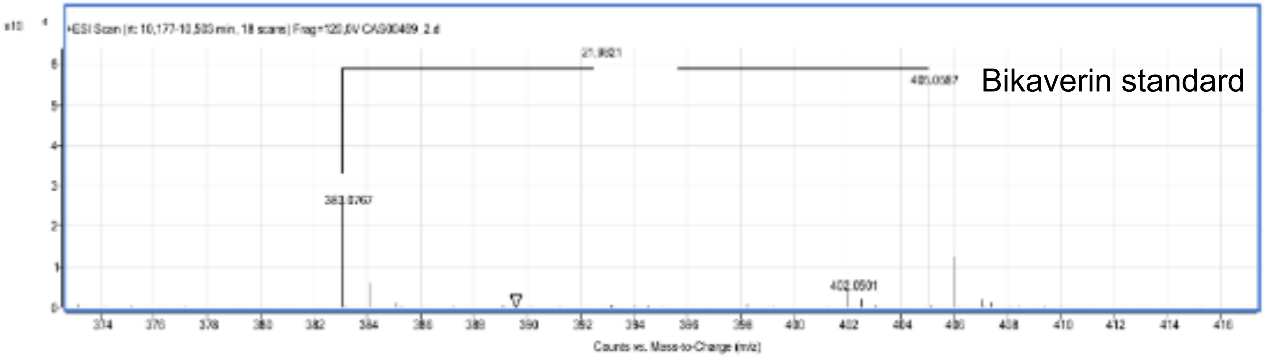

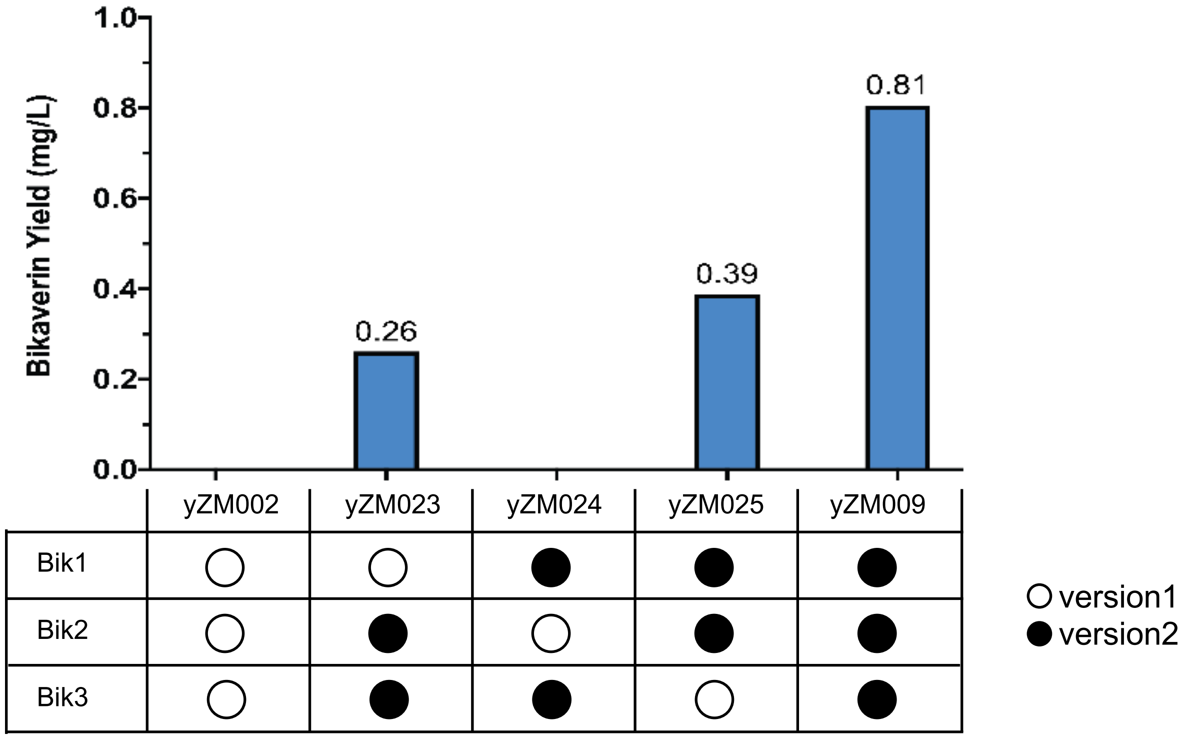


**Supplementary Fig. 3. Bikaverin production from V1 and V2 chimeric pathway.**

To figure out whether each V1-2007 enzyme is functional in yeast, we replaced each protein in functional bikaverin pathway from V2-2013 to V1-2007 version. Bikaverin was detected in yZM023, with Bik1 as V1 (at ~0.226mg/L) and yZM025, with Bik3 as V1 (at ~0.389mg/L), but both were lower than original strain yZM009 (at ~0.805mg/L). No bikaverin was detected in yZM024 with V1 Bik2 .These results indicated that in yeast, Bik2 enzyme with V1 amino acid sequence is not functional . The promoters of each gene are same as in S4a. Hollow circle stands version1 gene is in used, soild circle stands version 2 gene in used.

### HPLC/ESI-MS analytical standard

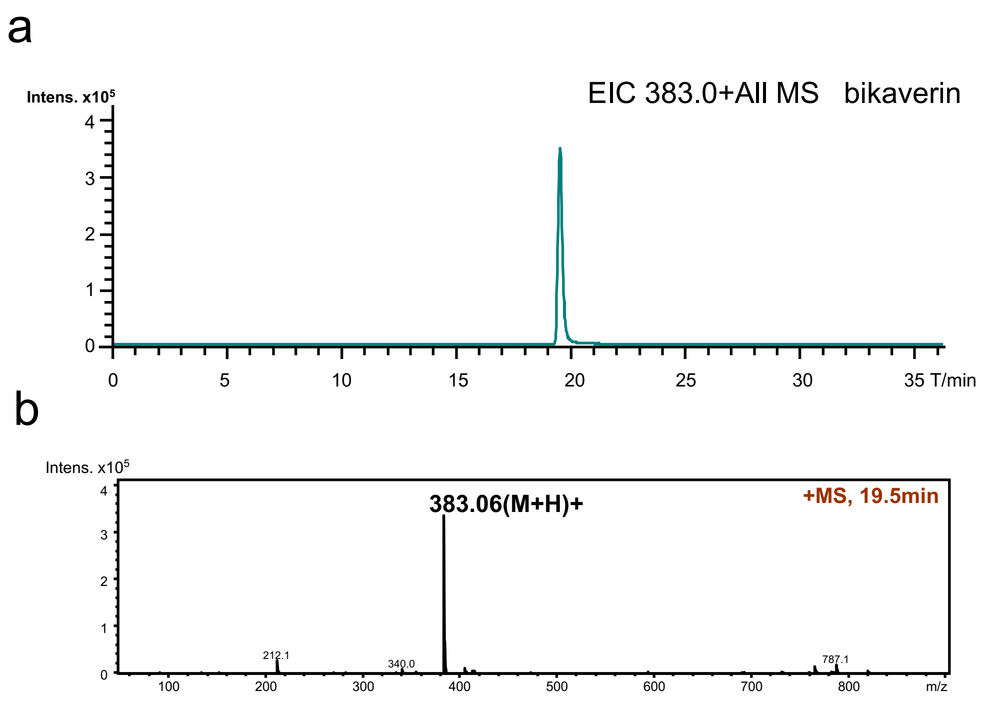


**Supplementary Fig. 4.** The Bikaverin standard for HPLC/ESI-MS analysis. **a.** the extracted ion chromatogram (EIC) of m/z 383.1**,** the mass of $[M\text{-}H]^+$ of bikaverin. **b.** the mass spectrum of the bikaverin peak in **a**.

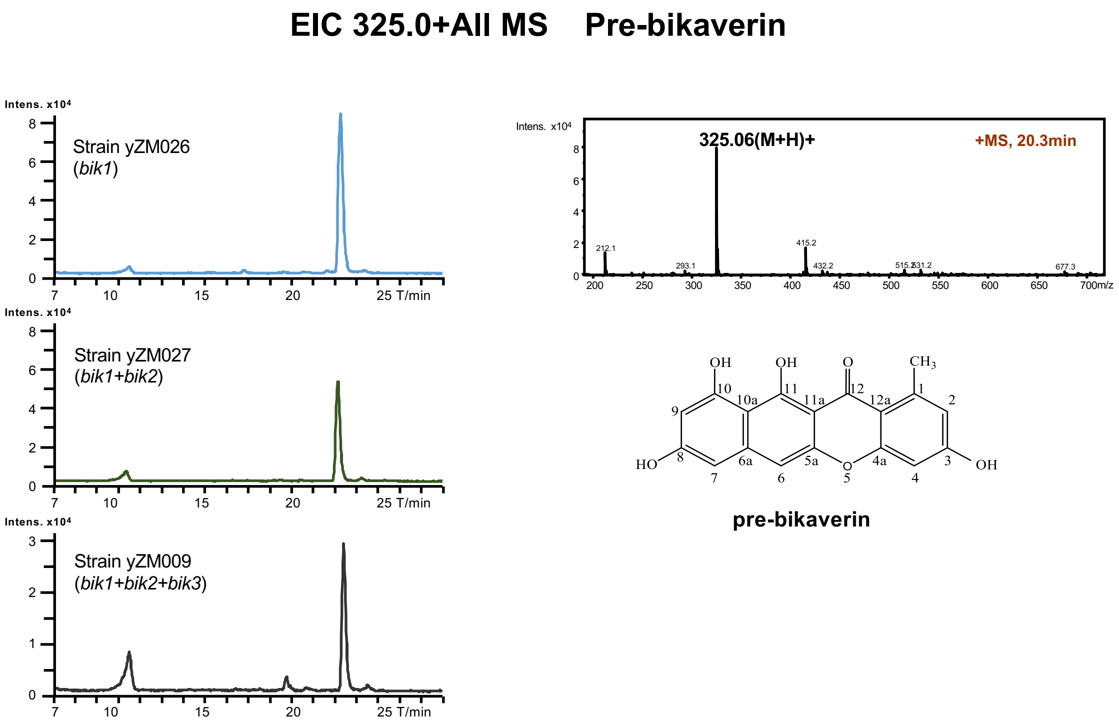


**Supplementary Fig. 5.** The EIC of pre-bikaberin with m/z 325.1 in strain yZM026, 027, 009. The Mass spectrum of the corresponding peak in yZM026 were shown in right side.

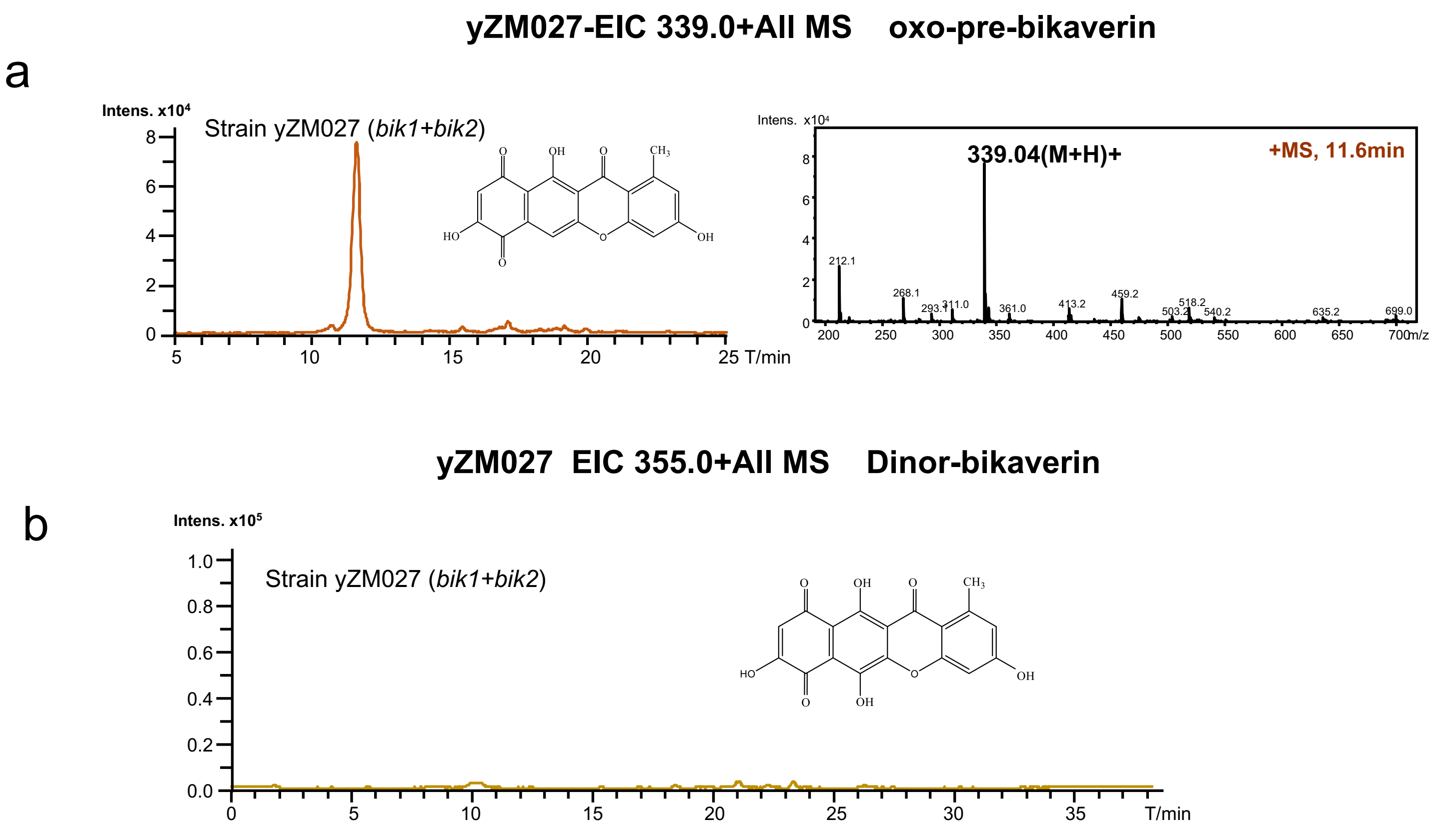


**Supplementary Fig. 6**. The EIC of oxo-pre-bikaberin and dinor-Bikaverin in strain yZM027. **a.** the peak of oxo-pre-bikaverin, and the corresponding mass spectrum. **b.** the extracted ion chromatogram of m/z 355.0, the mass of $[M\text{-}H]^+$ of dinor-bikaverin.

**yZM028-EIC 339.0+All MS    me-pre-bikaverin**

a

b

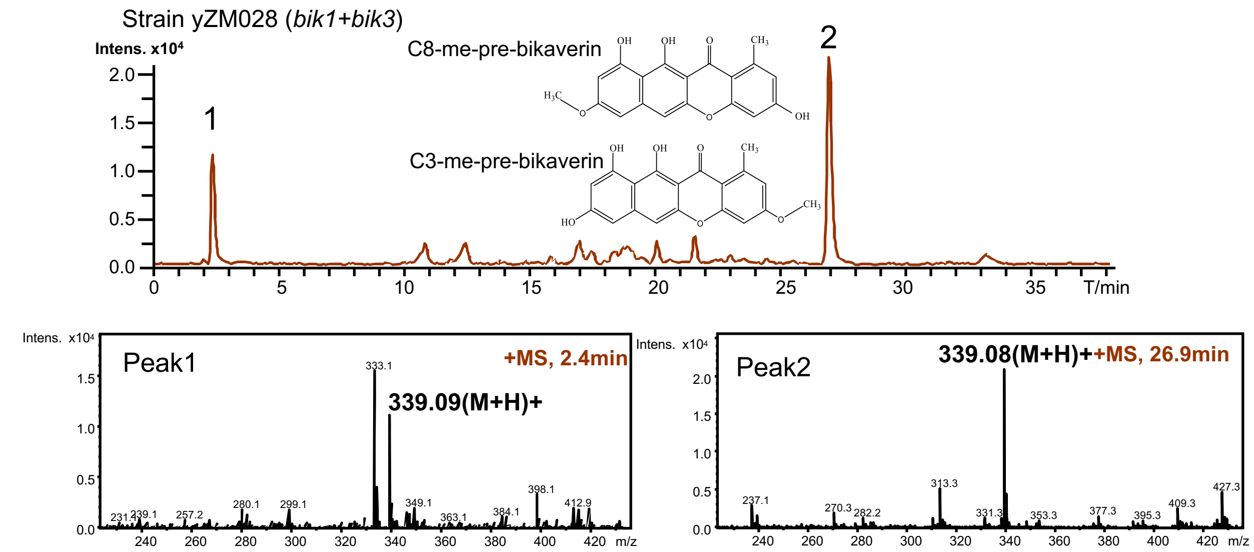


**Supplementary Fig. 7**

**a.** The two peak in EIC of me-pre-bikaberin (m/z 339 for $[M\text{-}H]^+$) from strain yZM028. **b.** The mass spectrum of the two peak were shown in below.

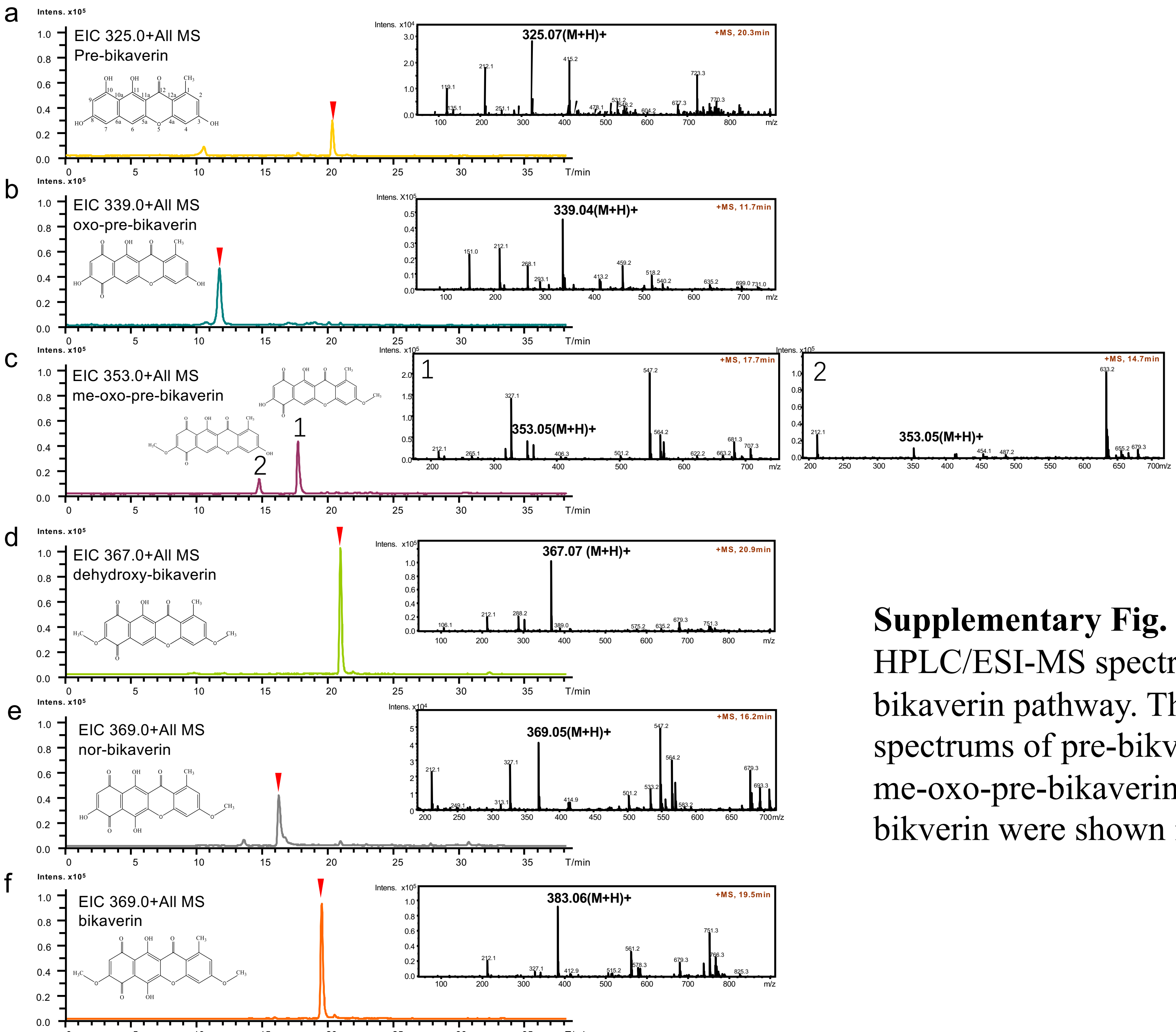


**Supplementary Fig. 8**

HPLC/ESI-MS spectrum of strain yZM009, with the complete bikaverin pathway. The extracted ion chromatograms and mass spectrums of pre-bikverin, oxo-pre-bikaverin, me-oxo-pre-bikaverin, dehydroxy-bikverin, nor-bikverin, bikverin were shown in **a**, **b**, **c**, **d**, **e** and **f**, respectively.

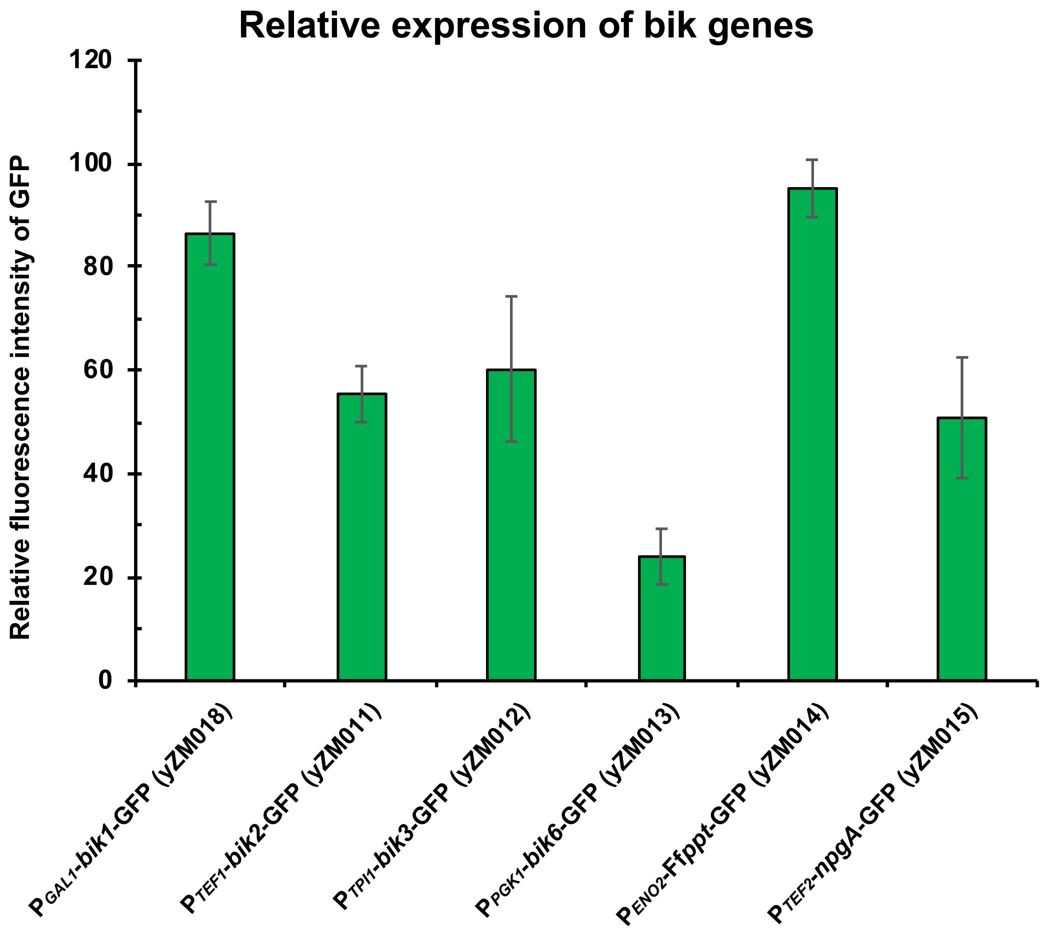


**Supplementary Fig. 9**

Relative expression of bik genes. All strains are grown in SC-Ura liquid medium with galactose as carbon source.

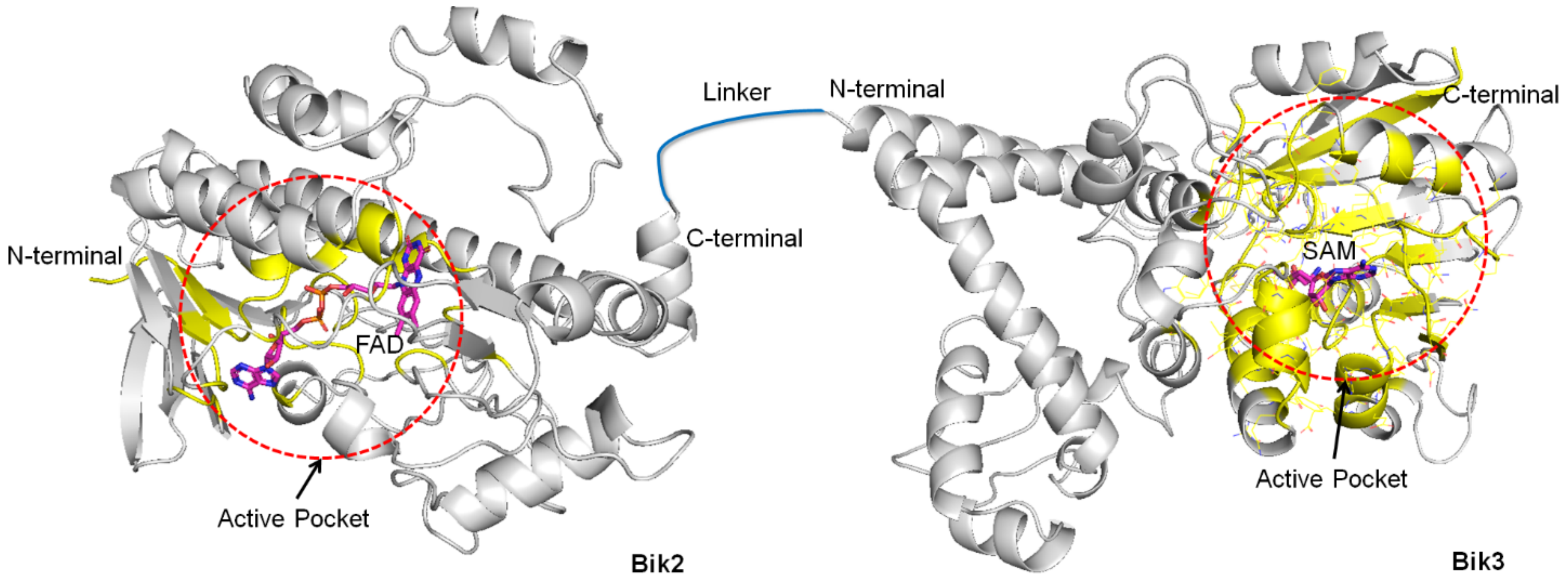


**Supplementary Fig. 10**

Bik2-Bik3 fusion design guided by homology modeling. The homology modeling shows that the active pocket and FAD bind site of Bik2 is near the N-terminal, while for bik3, its SAM binding site and active pocket are near the C-terminal. In order to not affect the functions of Bik2 and Bik3, we fusion the proteins in N terminal-Bik2-Bik3-C terminal direction.

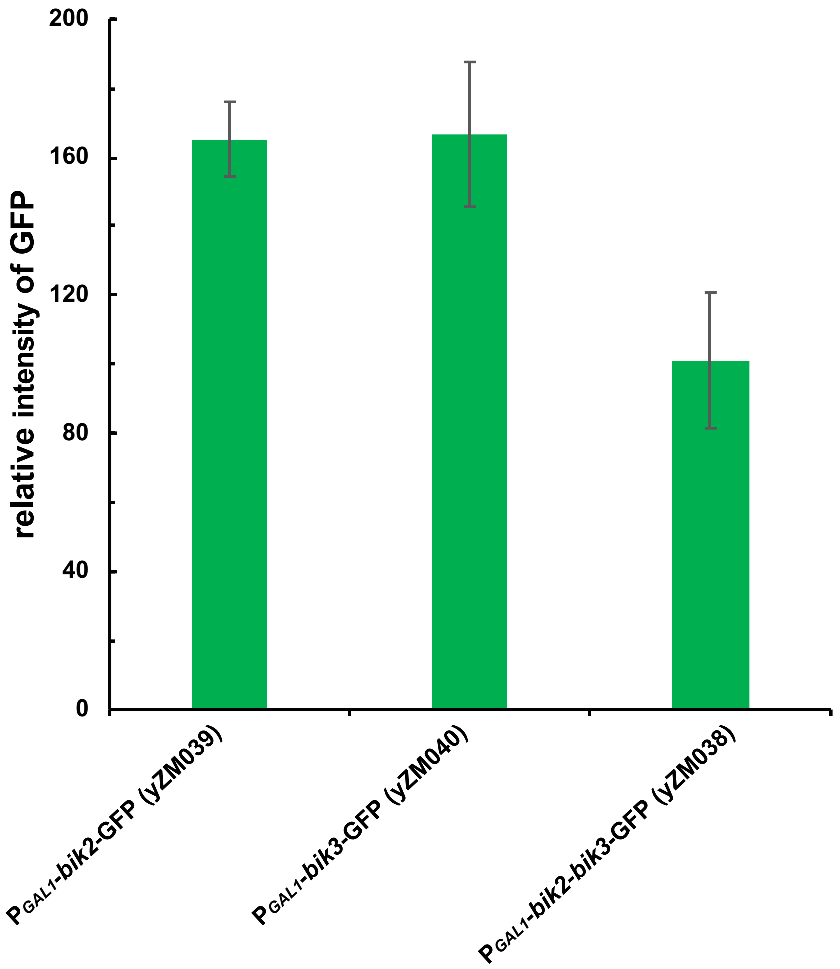


**Supplementary Fig. 11**
The expression level of Bik2-Bik3 fusion protein, compared to Bik2 and Bik3 expressed separately.
