## Supplementary Tables for "Pathway engineering in yeast for synthesizing the complex polyketide bikaverin"

This supplementary table word file includes:

**Supplementary Table 1.** Yeast strains used in this study.

**Supplementary Table 2.** E. coli strains and plasmids used in this study.

**Supplementary Table 3.** Primer sequence used for yeast homologous recombination.

**Supplementary Table 1.** Yeast strains used in this study.
Yeast strains and plasmids for expressing in yeast were listed in this table. Genes were marked with V1 meaning the Version 1genes. Unmarked genes were verison2 genes.

| Strain | Back-ground | Plasmids | Plasmids Description | |
|---|---|---|---|---|
| yZM001 | BY4742 | pZM-y001 | *pRS416_ PTEF1-BIK2(V1)- TCYC1_PTPI1-BIK3(V1)- TACS2_ PRPS2-BIK1(V1)- TACS1_PPGK1-BIK6(V1)- TZEO1_ PENO2-PPT-TADH2_PTEF2-NPGA- THXT7* | This study |
| yZM002 | BY4742 | pZM-y002 | *pRS416_ PTEF1-BIK2(V1)- TCYC1_PTPI1-BIK3(V1)- TACS2_ PGAL1-BIK1(V1)- TACS1_PPGK1-BIK6(V1)- TZEO1_ PENO2-PPT-TADH2_PTEF2-NPGA- THXT7* | This study |
| yZM006 | BY4742 | pZM-y006 | *pRS416_ PTEF1-BIK2- TCYC1_PTPI1-BIK3- TACS2_ PRPS2-BIK1-TACS1_PPGK1-BIK6- TZEO1_ PENO2-PPT- TADH2_PTEF2-NPGA- THXT7* | This study |
| yZM007 | BY4742 | pZM-y007 | *pRS416_ PTEF1-BIK2- TCYC1_PTPI1-BIK3- TACS2_ PRPL43A-BIK1-TACS1_PPGK1-BIK6- TZEO1_ PENO2-PPT- TADH2_PTEF2-NPGA- THXT7* | This study |
| yZM008 | BY4742 | pZM-y008 | *pRS416_ PTEF1-BIK2- TCYC1_PTPI1-BIK3- TACS2_ PGPM1-BIK1-TACS1_PPGK1-BIK6- TZEO1_ PENO2-PPT- TADH2_PTEF2-NPGA- THXT7* | This study |
| yZM009 | BY4742 | pZM-y009 | *pRS416_ PTEF1-BIK2- TCYC1_PTPI1-BIK3- TACS2_ PGAL1-BIK1-TACS1_PPGK1-BIK6- TZEO1_ PENO2-PPT- TADH2_PTEF2-NPGA- THXT7* | This study |
| yZM010 | BY4742 | pZM-y010 (Bik1-GFP) | *pRS416_ PTEF1-BIK2- TCYC1_PTPI1-BIK3- TACS2_ PRPS2-BIK1-GFP- TADH1_ PTEF-KanMX- TTEF- TACS1_PPGK1-BIK6- TZEO1_ PENO2-PPT- TADH2_PTEF2-NPGA- THXT7* ***(derien from pZM006）*** | This study |
| yZM011 | BY4742 | pZM-y011 (Bik2-GFP) | *pRS416_ PTEF1-BIK2-GFP- TADH1_ PTEF-KanMX- TTEF-TCYC1_PTPI1-BIK3- TACS2_ PRPS2-BIK1- TACS1_PPGK1-BIK6- TZEO1_ PENO2-PPT- TADH2_PTEF2-NPGA- THXT7* ***(derien from pZM006）*** | This study |
| yZM012 | BY4742 | pZM-y012 (Bik3-GFP) | *pRS416_ PTEF1-BIK2- TCYC1_PTPI1-BIK3-GFP- TADH1_ PTEF-KanMX- TTEF- TACS2_ PRPS2-BIK1- TACS1_PPGK1-BIK6- TZEO1_ PENO2-PPT- TADH2_PTEF2-NPGA- THXT7* ***(derien from pZM006）*** | This study |
| yZM013 | BY4742 | pZM-y013 (Bik6-GFP) | *pRS416_ PTEF1-BIK2- TCYC1_PTPI1-BIK3- TACS2_ PRPS2-BIK1-TACS1_PPGK1-BIK6-GFP- TADH1_ PTEF-KanMX- TTEF- TZEO1_ PENO2-PPT- TADH2_PTEF2-NPGA- THXT7* ***(derien from pZM006）*** | This study |
| yZM014 | BY4742 | pZM-y014 (PPT-GFP) | *pRS416_ PTEF1-BIK2- TCYC1_PTPI1-BIK3- TACS2_ PRPS2-BIK1-TACS1_PPGK1-BIK6- TZEO1_ PENO2-PPT-GFP- TADH1_ PTEF-KanMX- TTEF- TADH2_PTEF2-NPGA- THXT7* ***(derien from pZM006）*** | This study |
| yZM015 | BY4742 | pZM-y015 (NPGA-GFP) | *pRS416_ PTEF1-BIK2- TCYC1_PTPI1-BIK3- TACS2_ PRPS2-BIK1-TACS1_PPGK1-BIK6- TZEO1_ PENO2-PPT- TADH2_PTEF2-NPGA-GFP- TADH1_ PTEF-KanMX- TTEF- THXT7* ***(derien from pZM006）*** | This study |
| yZM016 | BY4742 | pZM-y016 (Bik1-GFP) | *pRS416_ PTEF1-BIK2- TCYC1_PTPI1-BIK3- TACS2_ PRPL43A-BIK1-GFP- TADH1_ PTEF-KanMX- TTEF- TACS1_PPGK1-BIK6- TZEO1_ PENO2-PPT- TADH2_PTEF2-NPGA- THXT7* ***(derien from pZM007）*** | This study |
| yZM017 | BY4742 | pZM-y017 | *pRS416_ PTEF1-BIK2- TCYC1_PTPI1-BIK3- TACS2_ PGPM1-BIK1-* | This |

| | | | | |
|---|---|---|---|---|
| | | (Bik1-GFP) | GFP- TADH1_ PTEF-KanMX- TTEF- TACS1_PPGK1-BIK6- TZEO1_ PENO2-PPT- TADH2_PTEF2-NPGA- THXT7 **(derien from pZM008)** | study |
| yZM018 | BY4742 | pZM-y018 (Bik1-GFP) | pRS416_ PTEF1-BIK2- TCYC1_PTPI1-BIK3- TACS2_ PGAL1-BIK1-GFP- TADH1_ PTEF-KanMX- TTEF- TACS1_PPGK1-BIK6- TZEO1_ PENO2-PPT- TADH2_PTEF2-NPGA- THXT7 **(derien from pZM009)** | This study |
| yZM021 | BY4742 | pZM-y006 +GEV | pRS413-GEV + pRS416_ PTEF1-BIK2- TCYC1_PTPI1-BIK3- TACS2_ PRPS2-BIK1- TACS1_PPGK1-BIK6- TZEO1_ PENO2-PPT- TADH2_PTEF2-NPGA- THXT7 | This study |
| yZM022 | BY4742 | pZM-y009 +GEV | pRS413-GEV + pRS416_ PTEF1-BIK2- TCYC1_PTPI1-BIK3- TACS2_ PGAL1-BIK1- TACS1_PPGK1-BIK6- TZEO1_ PENO2-PPT- TADH2_PTEF2-NPGA- THXT7 | This study |
| yZM023 | BY4742 | pZM-y023 | pRS416_ PTEF1-BIK2- TCYC1_PTPI1-BIK3- TACS2_ PGAL1-BIK1 (V1) - TACS1_PPGK1-BIK6- TZEO1_ PENO2-PPT- TADH2_PTEF2-NPGA- THXT7 | This study |
| yZM024 | BY4742 | pZM-y024 | pRS416_ PTEF1-BIK2 (V1) - TCYC1_PTPI1-BIK3- TACS2_ PGAL1-BIK1- TACS1_PPGK1-BIK6- TZEO1_ PENO2-PPT- TADH2_PTEF2-NPGA- THXT7 | This study |
| yZM025 | BY4742 | pZM-y025 | pRS416_ PTEF1-BIK2- TCYC1_PTPI1-BIK3 (V1) - TACS2_ PGAL1-BIK1- TACS1_PPGK1-BIK6- TZEO1_ PENO2-PPT- TADH2_PTEF2-NPGA- THXT7 | This study |
| yZM026 | BY4742 | pZM-y026 | pRS416_ PGAL1-BIK1-TACS1_PPGK1-BIK6- TZEO1_ PENO2-PPT-TADH2_PTEF2-NPGA- THXT7 | This study |
| yZM027 | BY4742 | pZM-y027 | pRS416_ PTEF1-BIK2- TCYC1 _PGAL1-BIK1-TACS1_PPGK1-BIK6-TZEO1_ PENO2-PPT- TADH2_PTEF2-NPGA- THXT7 | This study |
| yZM028 | BY4742 | pZM-y028 | pRS416_ PGAL1-BIK3- TACS2_ PGAL1-BIK1-TACS1_PPGK1-BIK6-TZEO1_ PENO2-PPT- TADH2_PTEF2-NPGA- THXT7 | This study |
| yZM029 | BY4742 | pZM-y029 | pRS416_ PGAL1-BIK2- TCYC1_PTPI1-BIK3- TACS2_ PGAL1-BIK1-TACS1_PPGK1-BIK6- TZEO1_ PENO2-PPT- TADH2_PTEF2-NPGA- THXT7 | This study |
| yZM030 | BY4742 | pZM-y030 | pRS416_ PTEF1-BIK2- TCYC1_PGAL1-BIK3- TACS2_ PGAL1-BIK1-TACS1_PPGK1-BIK6- TZEO1_ PENO2-PPT- TADH2_PTEF2-NPGA- THXT7 | This study |
| yZM031 | BY4742 | pZM-y031 | pRS416_ PGAL1-BIK2- TCYC1_PGAL1-BIK3- TACS2_ PGAL1-BIK1-TACS1_PPGK1-BIK6- TZEO1_ PENO2-PPT- TADH2_PTEF2-NPGA- THXT7 | This study |
| yZM032 | BY4742 | pZM-y032 | pRS416_ PTEF1-BIK2- TCYC1_PTPI1-BIK3- TACS2_ PGAL1-BIK1-TACS1_ PENO2-PPT- TADH2_PTEF2-NPGA- THXT7 | This study |
| yZM033 | BY4742 | pZM-y033 | pRS416_ PTEF1-BIK2- TCYC1_PGAL1-BIK3- TACS2_ PGAL1-BIK1-TACS1_ PENO2-PPT- TADH2_PTEF2-NPGA- THXT7 | This study |
| yZM034 | BY4742 | pZM-y034 | pRS416_ PGAL1-BIK2- TCYC1_PGAL1-BIK3- TACS2_ PGAL1-BIK1-TACS1_ PENO2-PPT- TADH2_PTEF2-NPGA- THXT7 | This study |
| yZM035 | BY4742 | pZM-y035 | pRS416_ PTEF1-BIK2- TCYC1_PTPI1-BIK3- TACS2_ PRPS2-BIK1-TACS1_PPGK1-BIK6- TZEO1_ PENO2-PPT- TADH2 | This study |
| yZM036 | BY4742 | pZM-y036 | pRS416_ PTEF1-BIK2- TCYC1_PTPI1-BIK3- TACS2_ PRPS2-BIK1-TACS1_PPGK1-BIK6- TZEO1_ PENO2-NPGA- TADH2 | This study |
| yZM037 | BY4742 | pZM-y037 | pRS416_ PGAL1-BIK2-BIK3- TCYC1_ PGAL1-BIK1-TACS1_ PPGK1-BIK6- TZEO1_PENO2-PPT- TADH2_PTEF2-NPGA- THXT7 | This study |

| | | | | |
|---|---|---|---|---|
| yZM038 | BY4742 | pZM-y038 | pRS416_ PGAL1-BIK2-BIK3-GFP- TADH1_ PTEF-KanMX- TTEF-TCYC1_ PGAL1-BIK1-TACS1_ PPGK1-BIK6- TZEO1_PENO2-PPT-TADH2_PTEF2-NPGA- THXT7 **（derien from pZM037）** | This study |
| yZM039 | BY4742 | pZM-y039 | pRS416_ PGAL1-BIK2-GFP- TADH1_ PTEF-KanMX- TTEF -TCYC1_PGAL1-BIK3- TACS2_ PGAL1-BIK1- GFP-TACS1_PPGK1-BIK6-TZEO1_ PENO2-PPT- TADH2_PTEF2-NPGA- THXT7 **（derien from pZM031）** | This study |
| yZM040 | BY4742 | pZM-y040 | pRS416_ PGAL1-BIK2- TCYC1_PGAL1-BIK3-GFP- TADH1_ PTEF-KanMX- TTEF - TACS2_ PGAL1-BIK1- GFP-TACS1_PPGK1-BIK6-TZEO1_ PENO2-PPT- TADH2_PTEF2-NPGA- THXT7 **（derien from pZM031）** | This study |

**Supplementary Table 2.** E. coli strains and plasmids used in this study.

| strain | Description | |
|---|---|---|
| bZM001 | *pUC19-VA1-PTEF1-BIK2 （V1） - TCYC1-VA3* | This study |
| bZM002 | *pUC19-VA3- PTPI1-BIK3 （V1） - TACS2-VA4* | This study |
| bZM003 | *pUC19-VA4- PRPS2-BIK1 （V1） - TAS1-VA5* | This study |
| bZM004 | *pUC19-VA5- PPGK1-BIK6 （V1） - TZEO1-VA6* | This study |
| bZM005 | *pUC19-VA6- PENO2-PPT （V1） - TADH2-VA7* | This study |
| bZM006 | *pUC19-VA7- PTEF2-NPGA （V1） - THXT7-VA2* | This study |
| bZM007 | *pUC19-VA1-PTEF1-BIK2- TCYC1-VA3* | This study |
| bZM008 | *pUC19-VA3- PTPI1-BIK3- TACS2-VA4* | This study |
| bZM009 | *pUC19-VA4- PRPS2-BIK1- TACS1-VA5* | This study |
| bZM010 | *pUC19-VA5- PPGK1-BIK6- TZEO1-VA6* | This study |
| bZM011 | *pUC19-VA6- PENO2-PPT- TADH2-VA7* | This study |
| bZM012 | *pUC19-VA7- PTEF2-NPGA- THXT7-VA2* | This study |
| bZM013 | *pUC19-VA6- PENO2-PPT- TADH2-VA2* | This study |
| bZM014 | *pUC19-VA6- PENO2-NPGA- TADH2-VA2* | This study |
| bZM015 | *pUC19-VA5- PENO2-PPT- TADH2-VA7* | This study |
| bZM016 | *pUC19-VA5- PENO2-PPT- TADH2-VA7* | This study |
| bZM017 | *pRS416-VA3-VA4- RFP-VA2* | |
| bZM018 | *pRS416-VA1- RFP-VA2* | |
| bZM019 | *pUC19-VA1-PGAL1-BIK2- TCYC1-VA3* | This study |
| bZM020 | *pUC19-VA3- PGAL1-BIK3- TACS2-VA4* | This study |
| bZM021 | *pUC19-VA4- PGAL1-BIK1- TACS1-VA5* | This study |
| bZM022 | *pUC19-VA4- PGAL1-BIK1 （V1） - TACS1-VA5* | This study |
| bZM023 | *pUC19-VA4- PRPL43A-BIK1- TACS1-VA5* | This study |
| bZM024 | *pUC19-VA4- PGPM1-BIK1- TACS1-VA5* | This study |
| bZM025 | *pUC19-VA4- PGPM1-BIK1 （V1） - TACS1-VA5* | This study |
| bZM026 | *pUC19-VA1-PGAL1-BIK2-BIK3 TCYC1-VA3* | This study |
| NA10 | NA10 | |
| GEV | GEV | |
| | pFA6a-GFP(S65T)-kanMX6 | |

**Supplementary Table 3.** Primer sequence used for yeast homologous recombination.

The sequence with underline were homologous arm used for yeast homologous recombination.

| Primer name | Sequence |
|---|---|
| Bik2-GFP-F | CCAGTTCAAGCTGCTACCGGTGTTGTTGAAGTTGGTTCTGGTCGACGGATCCCCGGGTT |
| Bik2-GFP-R | AACTAATTACATGATATCGACAAAGGAAAAGGGGCCTGTTCGATGAATTCGAGCTCGTT |
| Bik3-GFP-F | TACGGTTCTTTCATGTCTGTTATCGACGTTGTTTTGGGTGGTCGACGGATCCCCGGGTT |
| Bik3-GFP-R | CGAAATTTTATCTCATTACGAAATTTTTCTCATTTAAGTTCGATGAATTCGAGCTCGTT |
| Bik1-GFP-F | GCTTTGTGTGCTAAGATCAGAGAAACCATGGGTGTTAACGGTCGACGGATCCCCGGGTT |
| Bik1-GFP-R | AAAAAAAAGTCGTCAATATAAAAAGGAAAGAAATCATCATCGATGAATTCGAGCTCGTT |
| Bik6-GFP-F | ATCAGAGCTAGAGGTGAATTCTCTAAGTTGTCTACCTACGGTCGACGGATCCCCGGGTT |
| Bik6-GFP-R | AAAGAAACTTCTAGTAAAGTGCAGCACATTCAAGTGTGATCGATGAATTCGAGCTCGTT |
| PPT-GFP-F | GAAGAAATCTTGGCTTTCGGTGAACAAGCTTCTAAGCCAGGTCGACGGATCCCCGGGTT |
| PPT-GFP-R | AATGAAAACTATAAATCGTAAAGACATAAGAGATCCGCTTCGATGAATTCGAGCTCGTT |
| NPGA-GFP-F | ATCCAACCATGTGCTACCGGTGTTTGTAACTGTTTGTCTGGTCGACGGATCCCCGGGTT |
| NPGA-GFP-R | ATTAGAGCGTGATCATGAATTAATAAAAGTGTTCGCAAATCGATGAATTCGAGCTCGTT |
